# Benchmarking Cross-Docking Strategies for Structure-Informed Machine Learning in Kinase Drug Discovery

**DOI:** 10.1101/2023.09.11.557138

**Authors:** David Schaller, Clara D. Christ, John D. Chodera, Andrea Volkamer

## Abstract

In recent years machine learning has transformed many aspects of the drug discovery process including small molecule design for which the prediction of the bioactivity is an integral part. Leveraging structural information about the interactions between a small molecule and its protein target has great potential for downstream machine learning scoring approaches, but is fundamentally limited by the accuracy with which protein:ligand complex structures can be predicted in a reliable and automated fashion.

With the goal of finding practical approaches to generating useful kinase:inhibitor complex geometries for downstream machine learning scoring approaches, we present a kinase-centric docking benchmark assessing the performance of different classes of docking and pose selection strategies to assess how well experimentally observed binding modes are recapitulated in a realistic crossdocking scenario. The assembled benchmark data set focuses on the well-studied protein kinase family and comprises a subset of 589 protein structures co-crystallized with 423 ATP-competitive ligands. We find that the docking methods biased by the co-crystallized ligand—utilizing shape overlap with or without maximum common substructure matching—are more successful in recovering binding poses than standard physics-based docking alone. Also, docking into multiple structures significantly increases the chance to generate a low RMSD docking pose. Docking utilizing an approach that combines all three methods (Posit) into structures with the most similar co-crystallized ligands according to shape and electrostatics proofed to be the most efficient way to reproduce binding poses achieving a success rate of 66.9 % across all included systems.

The studied docking and pose selection strategies—which utilize the OpenEye Toolkit—were implemented into pipelines of the [KinoML framework](https://github.com/openkinome/kinoml) allowing automated and reliable protein:ligand complex generation for future downstream machine learning tasks. Although focused on protein kinases, we believe the general findings can also be transferred to other protein families.

## Introduction

Machine learning (ML) has found its way into many aspects of the drug discovery process [1]. Reliably predicting the bioactivity of small molecules binding to disease relevant targets has the potential to accelerate ligand design and to reduce costs especially in the early drug design phases [2]. While ligand-based ML models—which use the ligand chemical structure—are now widely used in drug discovery [3, 4, 5], structural information—where protein:ligand complex structures are encoded—in principle contains more information that can be used to improve affinity predictions. While it has so far proven difficult for structure-based methods to demonstrate practical improvements over ligand-based methods [6, 7, 8], the highly limited number of experimentally resolved protein:ligand complexes (compared to relatively more abundant protein:ligand affinity data) raised concerns about the generalizability of such models to a chemical space and protein targets not used in the training process [7, 9].

Several studies have assessed the ability of ML models utilizing protein:ligand complex structures to predict binding affinity data [7, 10, 11]. However, applying such approaches prospectively requires a reliable framework for generating experimentally unresolved protein:ligand complexes capable of capturing the relevant ligand binding mode(s) with sufficient accuracy. A number of new ML-based frameworks have been developed for predicting protein:ligand complexes that show potential [12, 13, 14], but have so far proven unable to rival traditional docking methods [15]. More traditional docking approaches sample docking poses inside a defined binding pocket of protein structures and select poses with physics-based scoring functions [16].

For structure-based machine learning methods to provide more utility than ligand-based methods alone, protein:ligand complex poses must be predicted with sufficient accuracy to both (1) allow poses and interactions to be determined prospectively for molecules that have not yet been synthesized, and (2) allow ligand poses to be inferred if experimental structures are unavailable for protein:ligand complexes for which affinity data is available and would be useful in model training. While re-docking benchmarks have proven useful in estimating the performance of docking tools to reproduce experimentally resolved X-ray structures [17, 18], the results cannot easily be translated into a prospective setting in which small molecules are docked into protein structures with binding pockets fitted to other co-crystallized ligands or without bound ligand. Furthermore, the strategy to select an experimentally resolved protein structure for docking a particular small molecule heavily influences the docking performance and is consequently equally important to assess. A few studies have been performed in this direction identifying the 3D similarity of the co-crystallized ligand and the binding site size as important parameters to select a suitable protein structure for docking [19, 20, 21]. However, these benchmark studies did not cover 2D similarity and were not embedded in fully automated pipelines, which is critical for high throughput complex generation and subsequent machine learning experiments on available binding affinity data.

The [OpenKinome](https://openkinome.org/) initiative represents a collaborative effort to build the infrastructure for controlled computational experiments towards structure-informed machine learning for bioactivity predictions of small molecules. For the development process the focus has been put on protein kinases, an important protein family for developing anticancer drugs [22]. Protein kinases are well-studied, resulting in abundant data available for both ligand bioactivity [23, 24] and experimentally resolved X-ray structures [25, 26], though with orders of magnitude more bioactivity measurements available than experimental structures. Here, we present the results of a cross-docking benchmark for docking pipelines implemented into the [KinoML](https://github.com/openkinome/kinoml) framework. We pay special attention to the conformational heterogeneity of protein kinases and evaluate several strategies to generate and select low RMSD docking poses. The code is made publicly available in the [kinasedocking-benchmark](https://github.com/openkinome/kinase-docking-benchmark) repository for the generation and analysis of the docking results, as well as in the [KinoML](https://github.com/openkinome/kinoml) repository for the corresponding docking pipelines.

## Results and Discussion

### Creation of a thorough cross-docking benchmark data set for protein kinases

The OpenCADD-KLIFS module [26, 27] was used to generate the cross-docking benchmarking data set for this study (**Table 1**). Prerequisite for kinase structures to be included in the benchmark data set was having a fully resolved ATP binding site with wild-type sequence to exclude any influence of modeling missing residues or mutations. Also, we included only kinases with at least 10 different structures of a kinase in a distinct KLIFS kinase conformation (namely DFG in/out and αC helix in/out, which correlate with the active and inactive states of a kinase [28]) with a single co-crystallized ligand in the ATP binding site. This procedure resulted in the final selection of 589 structures covering 10 distinct kinases and 423 ligands crystallized in a total of four different conformations (**Table 1, Appendix Figure 8**).

**Table 1.**
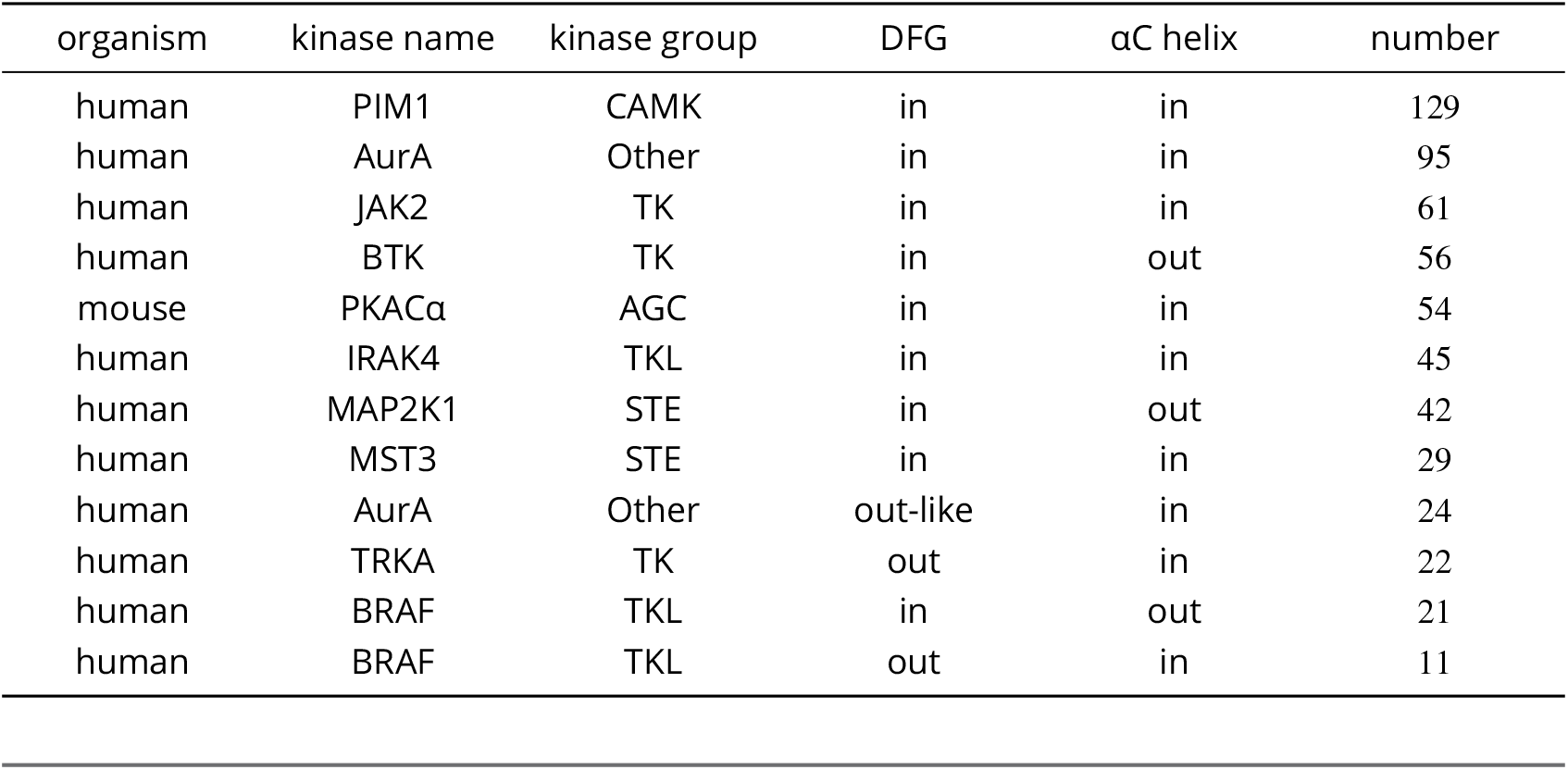
The kinase-inhibitor cross-docking benchmark set includes a representative set of kinases in a variety of conformations. A benchmark set of 589 kinase:inhibitor structures (representing 10 kinases in a variety of conformations) was constructed to ensure that ligands should be able to be robustly placed within the correct binding pose. We selected kinases with co-crystallized ligands in the ATP-binding site for which at least 10 structures were observed in the same DFG and αC helix conformation, and for which all 85 KLIFS residues contacting the ATP-binding site were both crystallographically resolved and contained wild-type residues only (benchmark set provided in **SI**).

Protein kinases are a conformationally heterogeneous protein family [29, 26]. ATP competitive inhibitors commonly prefer binding to a single kinase conformation, but have also found to bind to different (or multiple) kinase conformations depending on construct length, phosphorylation or mutations (**Table 2**) [30, 31, 32].

**Table 2.** Examples of orthosteric protein kinase ligands that bind to different kinase conformations. The KLIFS database was searched for orthosteric ligands binding to the same kinase in at least 3 different conformations. These observed conformations can result from alterations in the crystallized construct, e.g. construct length, co-crystallized nanobodies, mutations, and phosphorylation pattern. However, several ligands bind to different conformations of the same crystallized construct. Here, “Expo ID” refers to the chemical component identifier in the RCSB Ligand Expo [33] (or Chemical Components Dictionary).

| Expo ID | Kinase | DFG/αC helix conformation - PDB entry | Alterations |
| --- | --- | --- | --- |
| ATP | CAMK1 | in/in - 4FG7, in/out - 4FG9, out-like/out - 4FG8 | construct length |
| OWQ | MAPK6 | in/in - 6YKY (chain B), out-like/in 6YKY (chain A), out-like/out 6YKY (chain D) | none |
| OWX | MAPK6 | in/in - 6YLC (chain B), out-like/in 6YLC (chain A), out-like/out 6YLC (chain D) | none |
| ACP | AurA | in/in - 6CPF, out-like/in - 4C3R, out/in - 6C83 | nanobody |
| ADN | AurA | in/in - 4O0S, out-like/in - 4O0U, out/in - 1MUO | construct length, mutation |
| F9N | NEK7 | in/in - 6S73, in/out - 6S75 (chain B), out-like/out - 6S75 (chain A) | mutation |
| STU | MAP3K5 | in/in - 4BF2 (chain B), in/out - 4BF2 (chain A), out-like/in - 2CLQ (chain B), out-like/out - 2CLQ (chain A) | construct length mutation |
| B49 | MAK4K1 | in/in - 6NFZ, in/out - 6NG0 (chain A), out/out - 6NG0 (chain B) | phosphorylation, mutation |

**Table 3.** Benchmark data set creation. Sequential selection steps reduce the number of available KLIFS entries for docking from 12, 572 to 589.

| Selection step | Number of remaining KLIFS entries |
| --- | --- |
| All available KLIFS entries | 12,572 |
| Single orthosteric ligand | 10,439 |
| Highest quality chain per PDB structure | 5,017 |
| No alterations or unresolved residues in the KLIFS binding site | 1,942 |
| No ligands with unreasonable molecular weight or not handled by RDKit or OpenEye Toolkits | 1,915 |
| Exclude entries with multiple instances of the same ligand bound to the same protein chain | 1888 |
| At least 10 available structures per kinase and conformation from different kinase groups and conformations | 589 |

This observation complicates cross-docking studies across multiple kinase conformations, since a good docking score of an inhibitor to a kinase structure in a putative wrong conformation may not necessarily mean a poor performance of the docking algorithm. Instead this could also indicate that the inhibitor may actually also bind to this conformation, but was simply not yet resolved with a corresponding construct allowing it to crystallize in this conformation. Hence, this cross-docking study was only performed within the respective kinase conformations.

The stringent filtering criteria mentioned above reduce the total number of included structures for this docking benchmark tremendously and also exclude apo structures. In addition, this benchmark set is limited to a single protein class, i.e. protein kinases. However, the abundant structural data in conjunction with its detailed annotation in KLIFS [26] offers a unique chance to assess the docking performance in a realistic cross-docking scenario on an important drug target class. Also, we believe that the findings can be transferred to other target class with less available structural information.

### Molecular docking recovers most of the benchmark ligand poses

Three different docking algorithms were included in this study, all available via the OpenEye Toolkit [34]. Standard docking approaches sample possible conformations of flexible ligands in a rigid protein binding site and score the generated poses with a physics-based scoring function, i.e., **Fred**. In this study the binding site was defined by the 85 KLIFS residues accounting for the majority of protein:ligand interactions observed in X-ray structures of protein kinases [35]. The **Hybrid** method uses the shape and electrostatics of a co-crystallized small molecule as template to bias the placement of the new ligand in the binding site. The **Posit** method selects the most suitable docking algorithm based on the similarity of the co-crystallized ligand and the small molecule to dock, e.g., a small molecule differing only by a methyl substituent from the co-crystallized ligand will be placed via maximum common substructure docking [36], in contrast a small molecule with very low similarity will be docked with the physics-based method (Fred).

The cross-docking study was performed within structures of the same kinase and conformation (**Figure 1A**). For example, each of the 11 ligands co-crystallized with human BRAF in the DFG out/αC helix in conformation was docked into the remaining 10 structures of this kinase and conformation it was not co-crystallized with (**Table 1**). This procedure totals to ∼40K docking runs per crossdocking experiment, in which each ligand is docked into all other available structures of the kinase in the same conformation.

**Figure 1.**
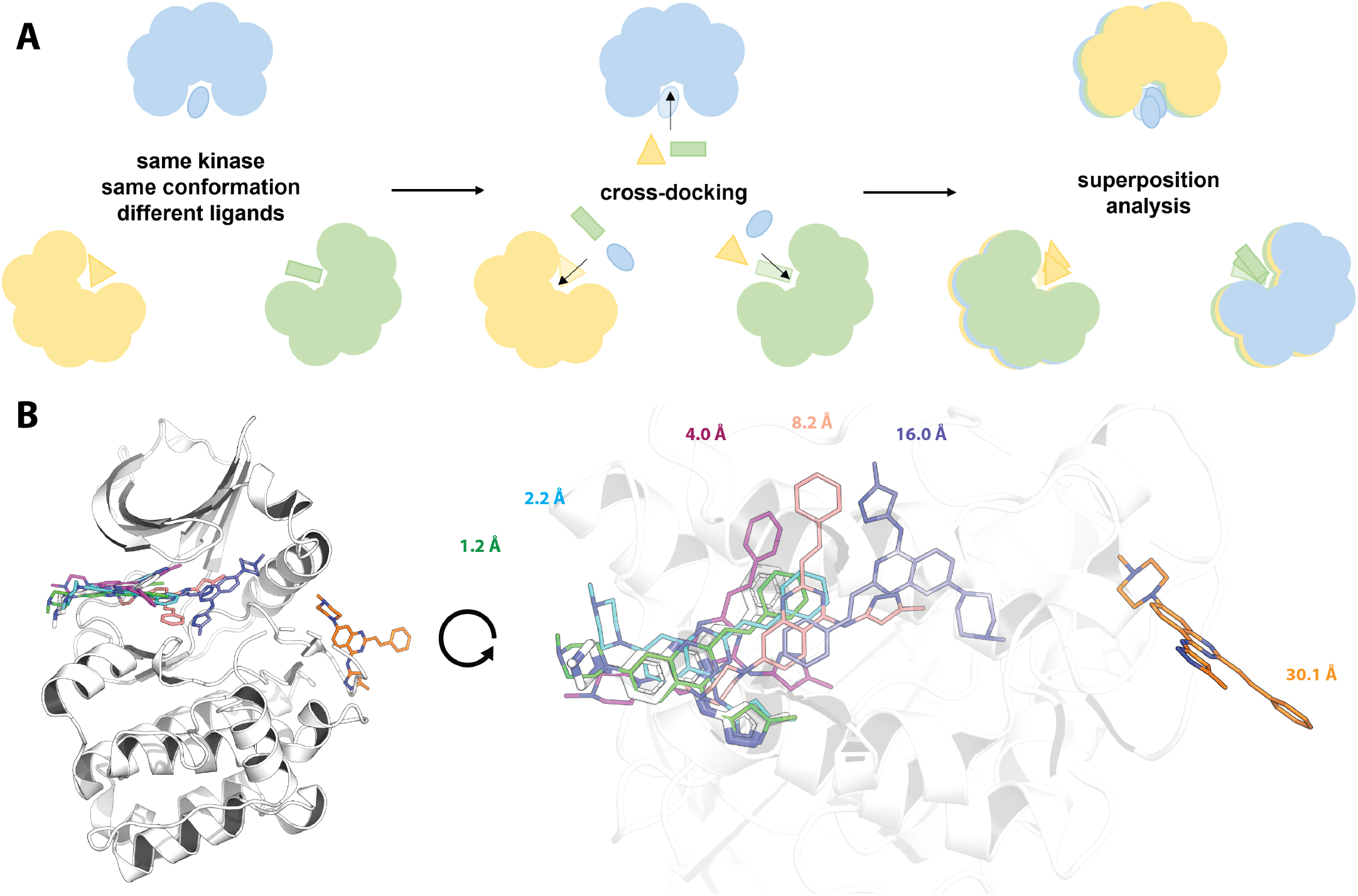
Design of this cross-docking study and visualization of different RMSD results observed in a cross-docking experiment. (A) Experimental design of this cross-docking study from structure selection, over the actual cross-docking experiment to the analysis after superposition. In the depicted example three X-ray structures (blue, yellow, green) of the same kinase resolved in the same conformation co-crystallized with three different ligands are used. Each ligand is docked into the orthosteric binding site of the other two kinase structures. The docked structures are then superimposed to the structure, in which the ligand was co-crystallized. Finally, the docking poses are analyzed via calculation of the root-mean-square deviation (RMSD). (B) The AurA entry 5ZAN with a co-crystallized ligand is depicted in white, docking poses generated by Fred for highlighted RMSD ranges in different colors. The docking pose with an RMSD below 2 Å recapitulates the binding mode and interactions observed in the X-ray structure. Docking poses with an RMSD up to 4 Å are located in the correct binding site. Docking poses with an RMSD of 8 Å or higher are located in other parts or outside of the binding pocket.

After docking, post-processing was performed with functionality from the OpenEye Toolkits [34] by superimposing the protein including the docking pose to the reference PDB structure with which the docked ligand was co-crystallized. Finally, the root-mean square deviation (RMSD) of heavy atoms between co-crystallized ligand and docking pose was calculated. A docking attempt generating a docking pose with an RMSD of ≤ 2 Å compared to the reference ligand structure was considered successful, which is a commonly used criterion in docking benchmark experiments [17, 18, 19, 20, 21]. Docking poses with an RMSD ≤ 4 Å usually cover the same binding pocket as the co-crystallized ligand, docking poses with an RMSD of ≥ 8 Å cover other parts of the binding pocket or are located outside the actual binding site (**Figure 1B**).

The RMSD analysis revealed that Posit was the most successful docking method achieving a mean bootstrapped success rate of 33.0 % when picking a random kinase structure of the same conformation for docking and 92.2 % when considering only the lowest RMSD pose per system to reproduce (**Figure 2** left, **Appendix Figure 9**). Physics-based docking alone (Fred) was the least successful method, reproducing the binding pose for only 23.8 % of systems when randomly selecting a kinase structure for docking and 84.3 % of systems in case of the lowest RMSD pose generate per system. These results highlight that the choice of a protein structure for docking a particular ligand has a dramatic impact on the performance, which was likewise shown by other studies before [19, 20, 21]. Additionally, the better performance of docking algorithms biased by a cocrystallized ligand is consistent with previous cross-docking benchmarks [19, 20].

**Figure 2.**
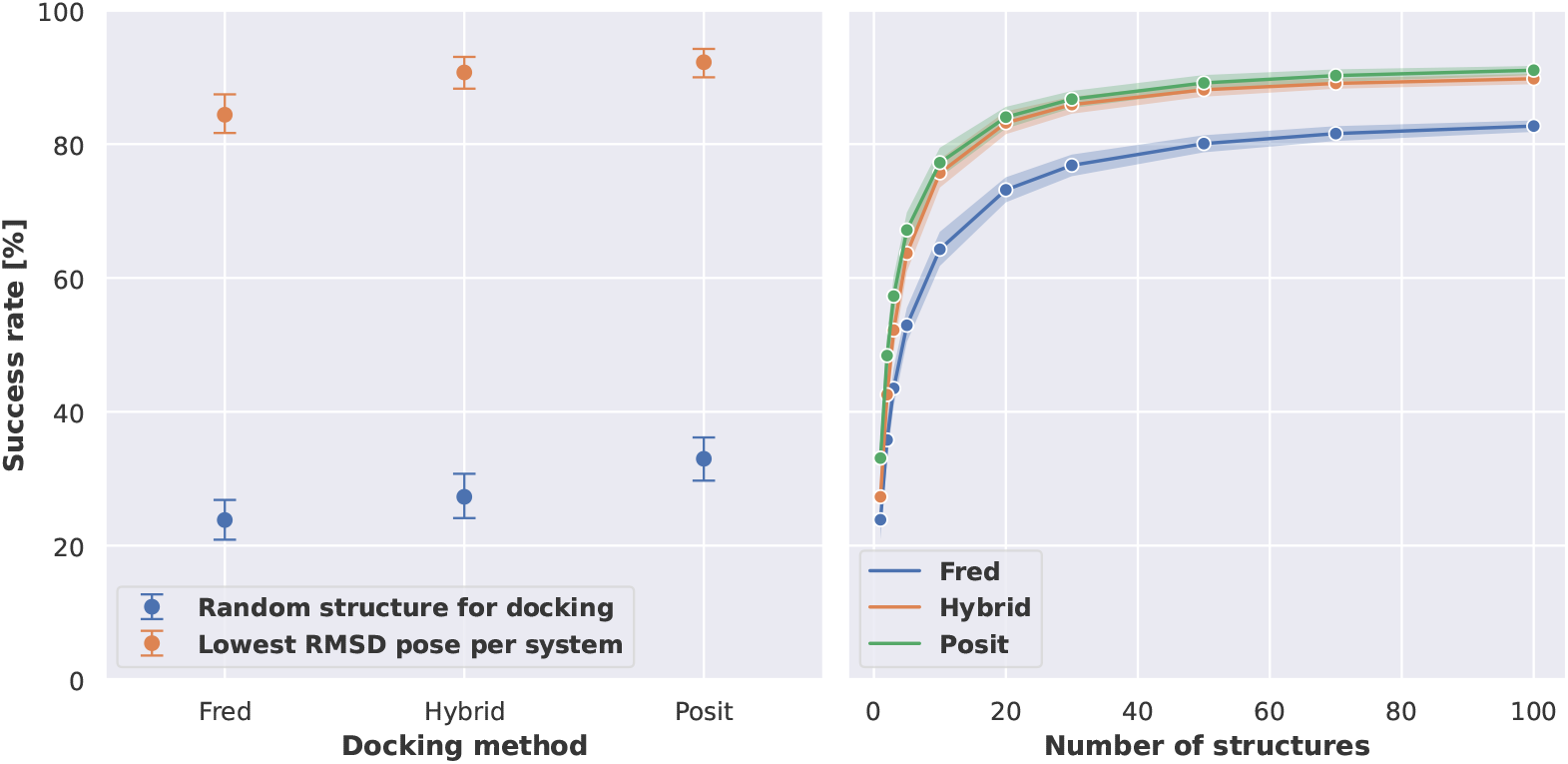
Molecular docking can recapitulate the majority of ligand binding poses but the success rate heavily depends on the selected kinase structure for docking. **Left:** Benchmark results show that low RMSD docking poses can be generated for the majority of the systems across the three docking method. The selection of the kinase structure for docking has a great impact on the success rate (pose ≤ 2 Å). **Right:** Success rates to generate docking poses ≤ 2 Å depend on the number of kinase structures used during docking. Observed differences between Hybrid and Posit are not statistically significant. Reported success rates and confidence intervals were estimated using bootstrapping.

### How can the chance be increased to generate and select a low RMSD docking Pose?

The success rates differ strongly when analyzing docking poses generated from docking into a randomly selected kinase structure compared to the lowest RMSD docking poses per system (**Figure 2** left). Hence, we were interested to identify strategies to increase the likelihood of generating and selecting low RMSD docking poses.

Due to the design of the benchmark data set, we could simulate the success rate of docking ligands into a different number of available X-ray structures and analyze the effect on generating a low RMSD docking pose. Therefore, the generated docking poses were bootstrapped to select docking poses for a defined number of randomly picked protein kinase structures. Increasing the number of protein structures used for docking steadily increased the chance to generate a low RMSD docking pose (**Figure 2** right). Posit and Hybrid methods required 20 structures to surpass a success rate of 80 %, Fred required 50 structures.

We also tested the effect of generating 5 instead of 1 pose per docking attempt (**Appendix Figure 11**). Generating multiple poses increased the success rate to generate a low RMSD pose when picking a random structure for docking. However, the success rate did not improve significantly when analyzing the lowest RMSD pose generated per system. Adding the results for docking into multiple structures (**Figure 2**), one can conclude that docking into multiple kinase structures is more successful than generating multiple docking poses for a single kinase structure.

Previous studies have found a positive effect of docking into structures with similar co-crystallized ligands [19, 20]. We could observe a similar behavior with regard to 2D similarity using Morgan fingerprints and 3D similarity estimated with shape and electrostatic criteria (**Figure 3**). However, restricting docking runs to structures with more similar co-crystallized ligands, led to the exclusion of many systems for which no co-crystallized ligand was available meeting the required similarity threshold. For example applying a 2D similarity cutoff of 0.8 increased the success rate for Posit from 33.0 % to almost 80 %. However, at this cutoff only 40 % of the systems had a corresponding kinase structure available.

**Figure 3.**
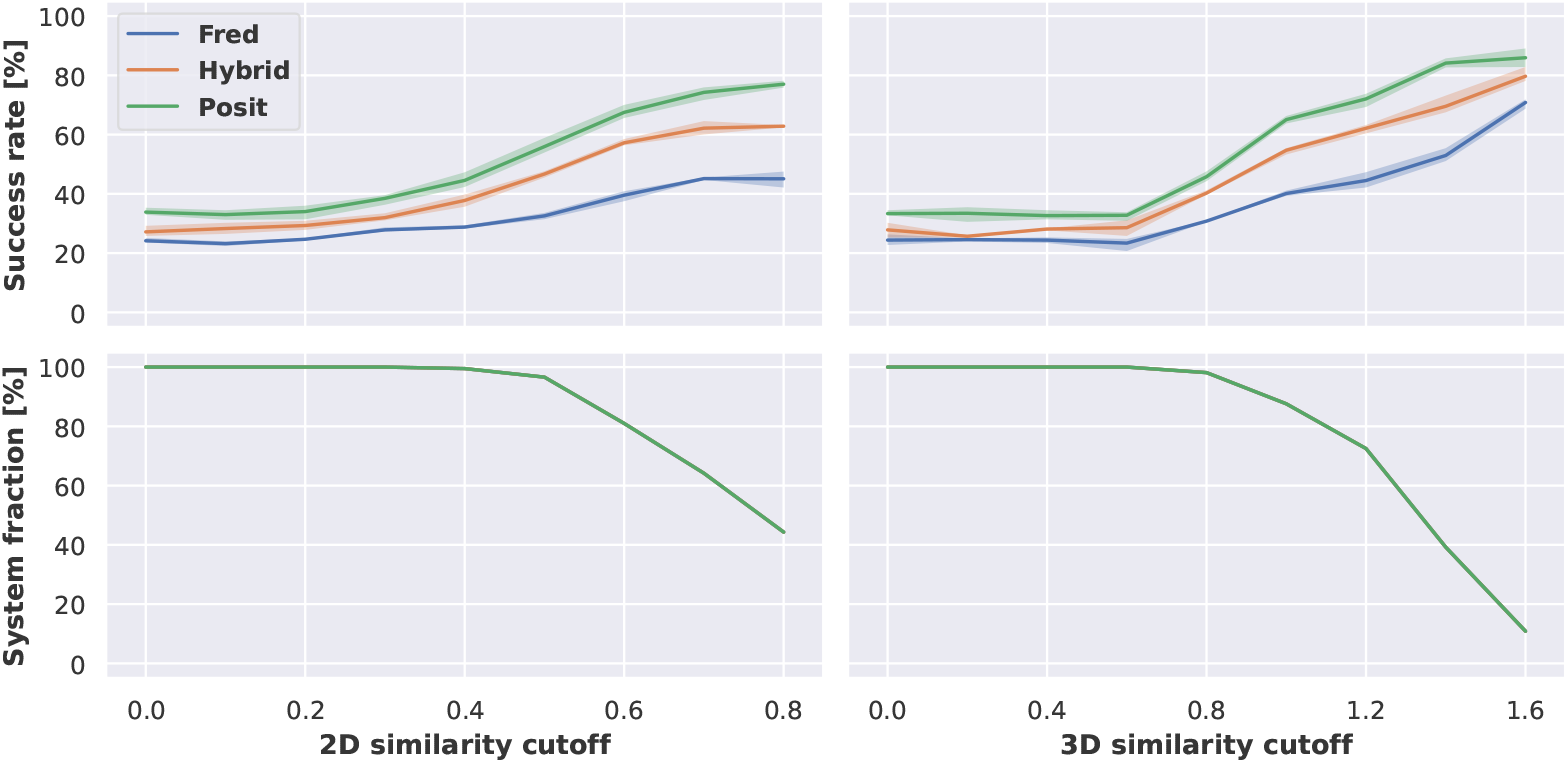
Docking success rates improve when docking into structures with more similar co-crystallized ligands. The 2D similarity of ligands was assessed using Morgan fingerprints via Dice similarity implemented in [RDKit](http://www.rdkit.org/), 3D similarity using overlay of shape and electrostatics via Tanimoto similarity implemented in OpenEye Toolkits [34]. Reported success rates and confidence intervals were estimated using bootstrapping.

Given the increased availability of computational resources, it would be also a viable strategy to dock into all available kinase structures and afterwards restrict the analysis to docking poses with sufficiently good docking scores. In this study, we analyzed the Chemgauss4 docking score for all three docking methods and additionally the Posit probability for the Posit method. The Posit probability estimates the likelyhood of generating a low RMSD pose based on the 2D and 3D similarity of the ligand to dock and the co-crystallized small molecule (see the [documentation](https://docs.eyesopen.com/applications/oedocking/theory/posit_theory.html#posit-heat-map) for further details). The docking score was normalized by the heavy atom count of the docked molecule to account for the correlation between docking score and molecular weight [37].

We observed an increasing overall success rate when applying a more stringent docking score cutoff (**Figure 4**). This trend reverts for Hybrid and Fred when surpassing a heavy atom count normalized docking score cutoff of −0.8. However, there are less than 20 % of the systems left at this cutoff, which is also reflected in an increased confidence interval. Additionally, we observed a relatively large fraction of ATP analogues in the remaining systems at such high cutoffs, which may add an artificial bias to the docking performance. For example the phosphate groups of the ATP analogues have multiple charged centers which can perform strong charged interactions and consequently result in good docking scores. Contrary, the phosphates of the ATP analogues are very flexible and thus might be relatively hard to position correctly for a low RMSD docking pose. Applying more stringent Posit probability cutoffs also excluded several systems from analysis, for which no sufficient docking pose could be generated. But in comparison to the normalized docking score this affected far less systems. For example at a Posit probability cutoff of 0.9, which is very close to theoretical maximum of 1, 50 % of the systems remained for analysis and showed a success rate of over 80 % in generating low RMSD docking poses.

**Figure 4.**
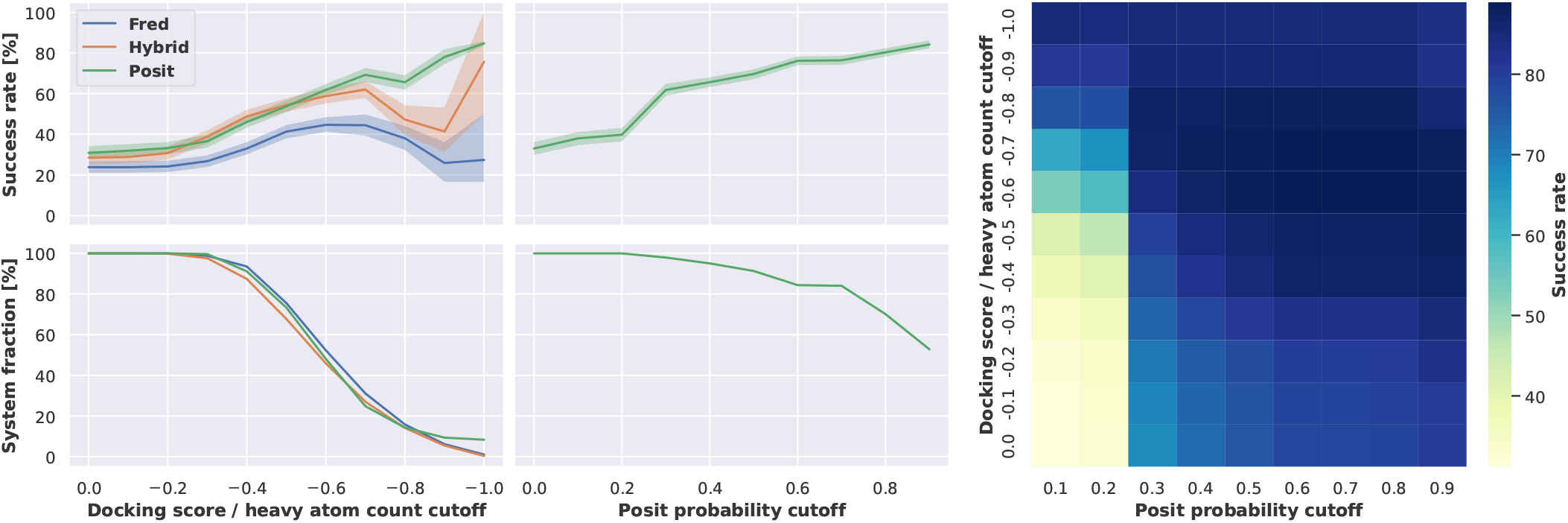
Docking success rates improve when applying more stringent docking score or Posit probability cutoffs at the expense of suitable systems for docking. The docking success rate increases to more than 60 % for Fred and Hybrid when only picking from docking poses with a heavy atom count normalized docking score equal to or lower than −0.6. However, this also excludes 50 % of the benchmark structures, for which no docking poses were generated satisfying this cutoff. In contrast, a Posit probability cutoff of 0.3 also allows for a docking success rate of 60 % but excludes only less than 5 % of the benchmark systems. Reported success rates and confidence intervals were estimated using bootstrapping. The heat map shows that docking poses with a Posit probability below 0.3 can have a docking success rate of more than 60 % if the heavy atom count normalized docking score is −0.8 or lower. Another version of this figure without normalization of the docking score is available in the appendix (**Figure 10**), but the overall findings do not change.

Overall, the Posit probability was found to be more successful in selecting low RMSD docking poses than the normalized docking score. However, the normalized docking score was found especially valuable for docking poses with a Posit probability below 0.3 (**Figure 4 right**). Here, a normalized docking score below −0.7 could reliably predict low RMSD docking poses. This finding suggests, that both metrics could also be combined to pick low RMSD docking poses.

Next, we compared different docking methods and docking pose selection strategies on the full benchmark data set without applying any similarity or docking score cutoffs (**Figure 5**). Two extremes were included to help interpreting the success rates. The **random** strategy randomly selects a kinase structure for docking for each system. In contrast, the **ideal** strategy reflects a situation, in which we know which structure will deliver the lowest RMSD docking pose for each structure that is being reproduced. Both extremes show the expected range of success rates for different strategies, for which we should not perform worse than the random strategy but will also not perform better than the ideal strategy. The best combination of docking method and selection strategy was the Posit algorithm selecting the best docking pose according to the Posit probability (73.3 % success rate). However, this was not statistically significantly better than picking the X-ray structure for docking with the most similar co-crystallized ligand according to 3D similarity reflecting shape and electrostatics (66.9 % success rate using 3D ligand similarity). This is important because selecting a structure based on Posit probability requires docking into all structures, while the 3D similarity to the co-crystallized can be calculated beforehand. A promising option is available in the OpenEye Toolkits [34], in which the user can provide multiple protein structures for docking with Posit and an automated routine will pick the most suitable protein structure for docking based on 2D and 3D similarity between co-crystallized ligand and small molecule to dock. This option was not included in this study, since we were interested into receiving docking poses for all structures.

**Figure 5.**
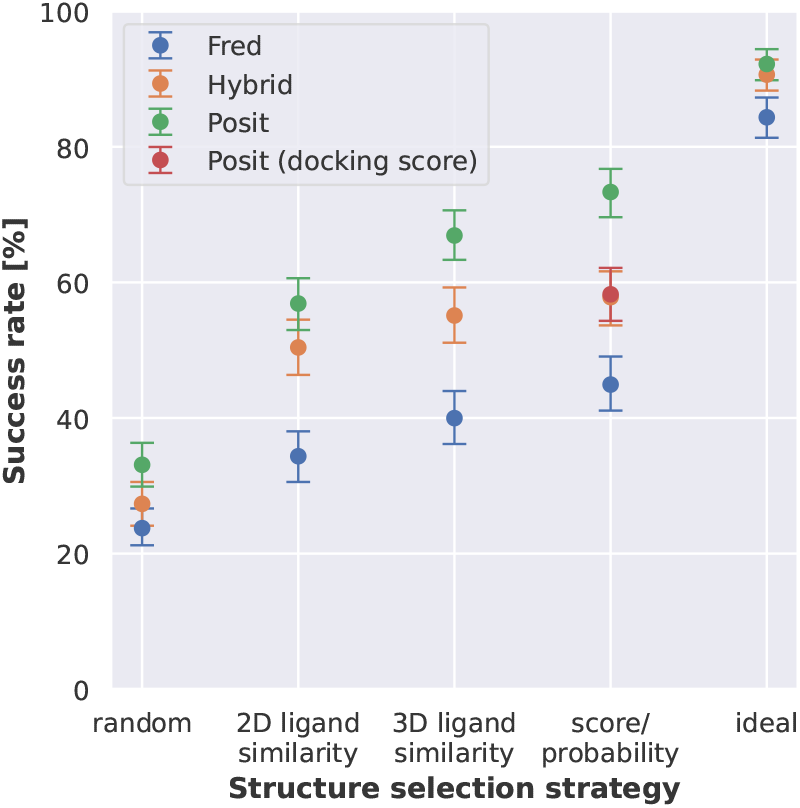
Kinase inhibitor pose recovery in cross-docking benefits from docking to multiple structures, selection of optimal ligand templates, and leveraging ligand information in an adaptive fashion (Posit). Posit clearly outperforms Hybrid and Fred. Success rates (pose below 2 Å) over 589 kinase:inhibitor pairs. Kinase structure for docking randomly selected, by 2D or 3D ligand similarity to co-crystallized ligand, by docking score / Posit probability, or by identifying the lowest RMSD pose (ideal scenario). Reported success rates and confidence intervals were estimated using bootstrapping.

Overall, Posit was found to be the most successful docking method in reproducing binding poses from X-ray structures (**Figures 2** and **5**). However, Posit also generated more poses with a higher (more unfavorable) docking score than Fred and Hybrid (**Appendix Figure 12**), which indicates steric clashes or high strain ligand conformations. This is likely caused by a strong bias from the co-crystallized ligand when placing the new ligand in the binding site. An energy minimization or short molecular dynamics simulation could potentially relax such poses. A pose relaxation option is available in the OpenEye Toolkits [34], but this has not been tested in this study, since we wanted to focus this study on the docking performance and not different force field behaviors.

### Transferring ligands for Posit docking does not improve performance

After discovering the good performance of Posit, we implemented and tested a customized docking protocol employing Posit with transferred co-crystallized ligand template. In this protocol, the KLIFS database is searched for similar co-crystallized ligands according to shape and electrostatics compared to the ligand that should be docked. The search is thereby covering all available kinase structures limited to the same DFG and αC helix conformation. The structure with the most similar ligand is then superimposed using functionality from the OpenEye Toolkits [34] to the structure the ligand should be docked into and the new more similar ligand is transferred. This procedure was thought to improve the performance of the Posit method, since it is highly dependent on the similarity to the co-crystallized ligand (**Figure 3**).

As expected, the similarity of the ligand template used to bias the Posit method could be increased by transferring more similar ligands from other kinase structures (**Appendix Figure 13**). The success rate for randomly picked kinase structures for docking increases from 33.0 % to 47.3 % when transferring a more similar template ligand from another kinase structure (**Figure 6**). However, the success rate for the lowest RMSD pose generated per system is decreasing from 92.2 % to 54.8 %. Additionally, the success rate of the lowest RMSD pose per system (54.8 %) is worse than docking into structures with the most similar co-crystallized ligand according to 3D similarity without applying the transfer protocol (66.9 %, **Figure 5**). This shows that transferring more similar ligands from other kinases does not always improve docking performance. However, ligand transfer was also performed for very distantly related kinases, e.g. ligands from atypical kinases where transferred into structures of tyrosine kinases. Restricting the ligand transfer to be performed only with a kinase group may improve the performance but was not tested in this study.

**Figure 6.**
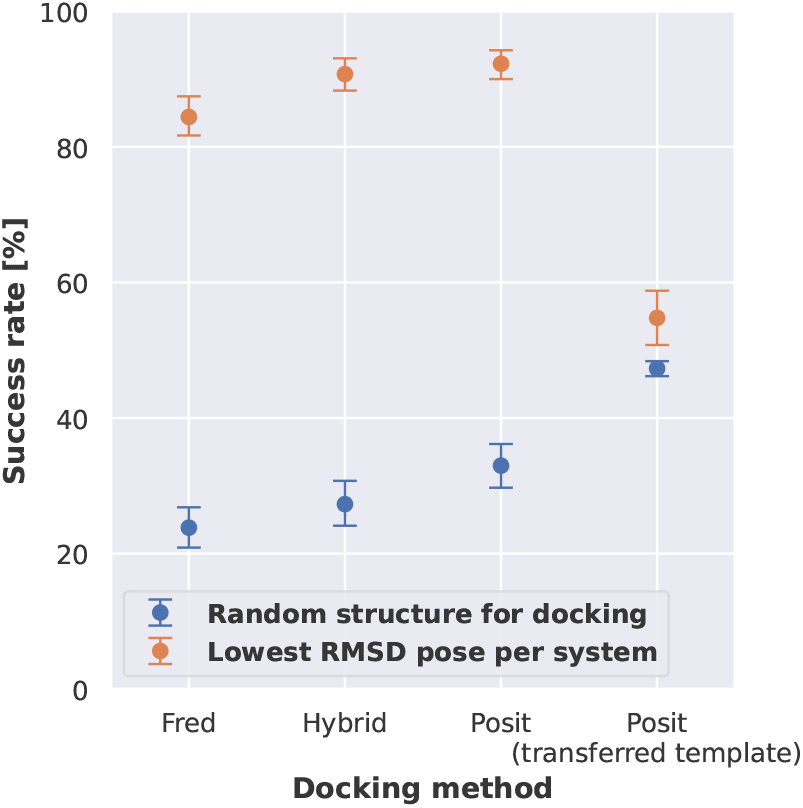
Transferring more similar ligands from other kinase structures improves the success rate for Posit when docking into a single kinase structure but not when docking into multiple kinase structures. A customized docking protocol was implemented employing Posit with a transferred co-crystallized ligand template. The ligand co-crystallized to each structure was thereby replaced by the most similar ligand compared to the ligand to dock found in the KLIFS database according to shape and electrostatics.

## The Posit probability can discriminate different kinase conformations with larger structural rearrangements

The benchmark data set contains two kinases crystallized in different activation loop conformations, i.e., human AurA and human BRAF (**Table 1**). To assess whether docking strategies could also identify the correct kinase conformation, another experiment was performed docking each ligand into all other structures of the kinase it was not co-crystallized with including a different DFG and αC helix conformation. Here, we wanted to assess (i) if the most successful docking strategy, Posit with picking a docking pose via posit probability (**Figure 5**), can identify the kinase conformation it was co-crystallized with and (ii) if including multiple conformations has an effect on the overall docking performance.

The conformational changes for both AurA and BRAF were first analyzed in terms of Pocket RMSD calculated for the 85 KLIFS binding site residues (**Figure 7** right) [35]. According to the RMSD the conformation change observed in case of BRAF (out/in vs. in/out) is larger compared to AurA (in/in out-like/in).

**Figure 7.**
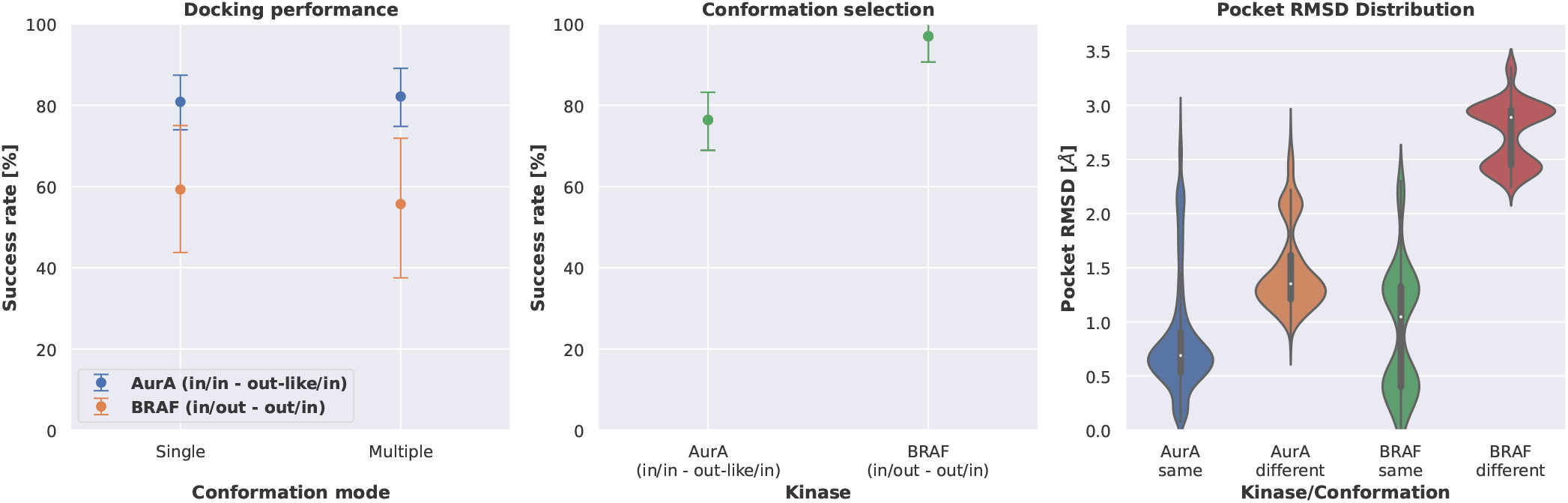
The Posit docking performance is not significantly affected by docking into kinase structures of different conformations. The Posit probability can pick the correct kinase conformation in the majority of included structures. The docking benchmark set contains structures for two kinases in two different conformations (DFG/αC helix motifs), i.e. AurA: in/in vs out-like/in and BRAF: out/in vs in/out (**Table 1**). Left: The docking success rate is not statistically different when selecting the docking pose with the best Posit probability from docking poses generated by docking into structures with the correct conformation or with two different conformations. Middle: Detecting the correct kinase conformation with the Posit probability is more successful for larger structural differences, i.e. out/in vs in/out compared to in/in vs out-like/in. Right: The larger structural rearrangement in case of BRAF (in/out vs. out/in) compared to AurA (in/in vs. out-like/in) is reflected in the pocket RMSD, which rationalizes the better conformation selection success rate for the BRAF kinase.

**Figure 8.**
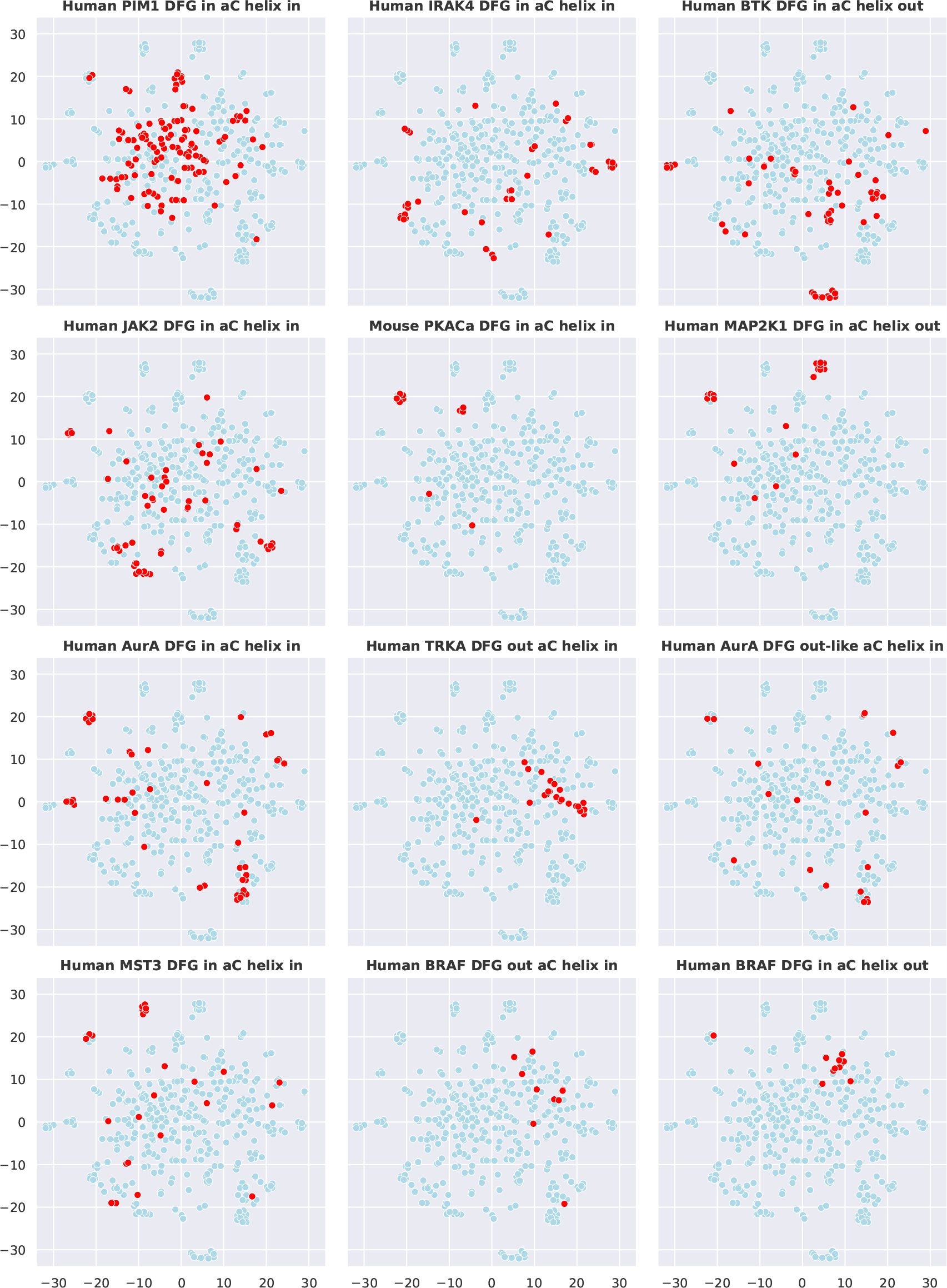
Chemical space of co-crystallized ligands for each kinase and conformation included in the docking benchmark data set. The chemical space is visualized via T-SNE representation of 50 principal components accounting for 64 % of observed variance in the principal component analysis. Blue dots represent ligand data points of the entire data set, red dots of the respective kinase and conformation. The principal components are based on Morgan fingerprint with features of radius 2 and 2, 048 bits implemented in the [RDKit](http://www.rdkit.org/). The principal components analysis and T-SNE representation was performed with Scikit-learn [41].

**Figure 9.**
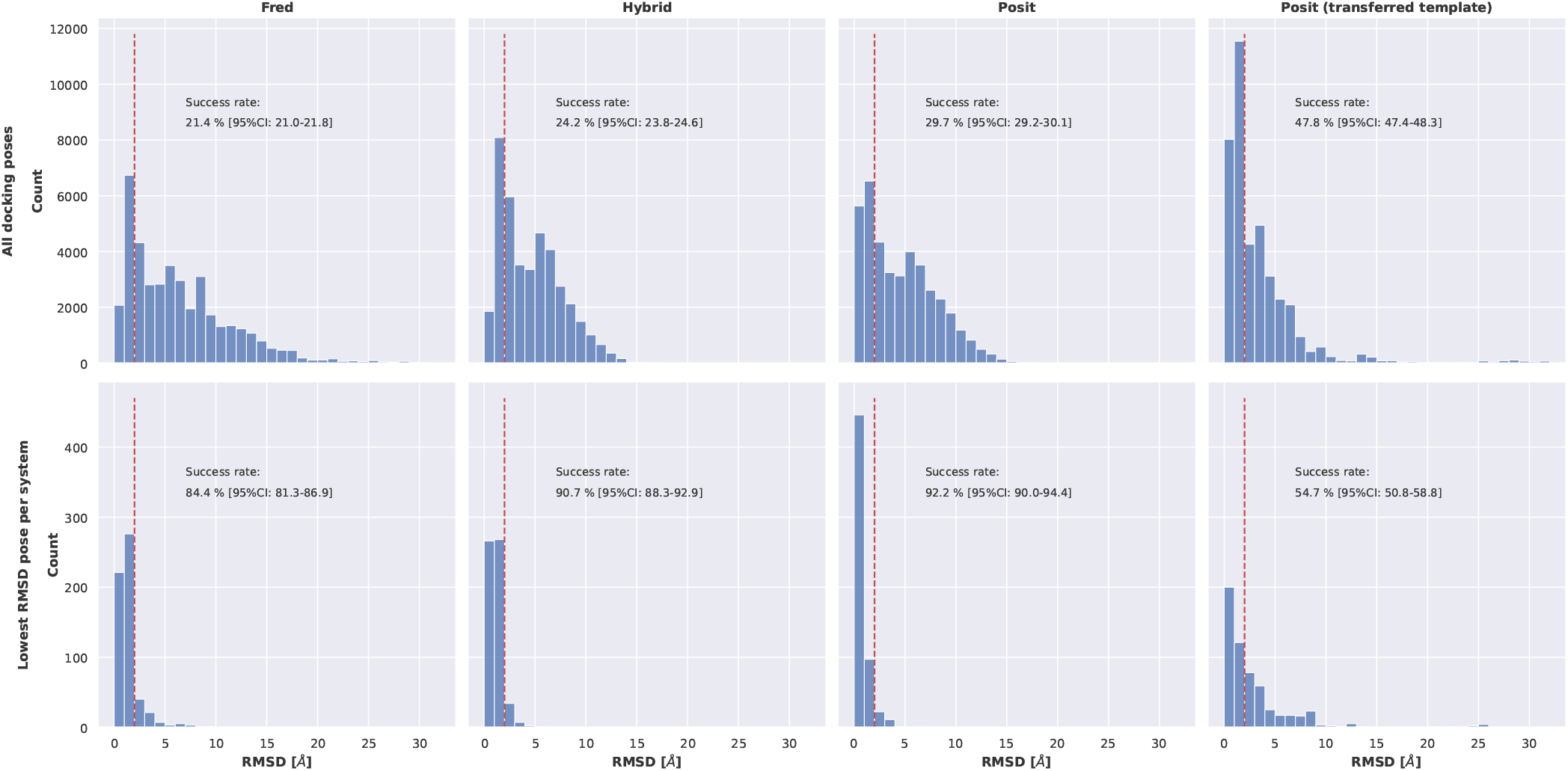
Assessment of the accuracy of kinase inhibitor pose recovery from cross-docking into kinase structures in the same conformation. Benchmark results show that low RMSD docking poses can be generated for the majority of the systems; docking into multiple structures is important. Top row: Average success rates (pose below 2 Å) for all ∼40K docking poses. Bottom row: Lowest RMSD poses for 589 kinase:inhibitor pairs in total. Reported success rates and confidence intervals were estimated using bootstrapping.

**Figure 10.**
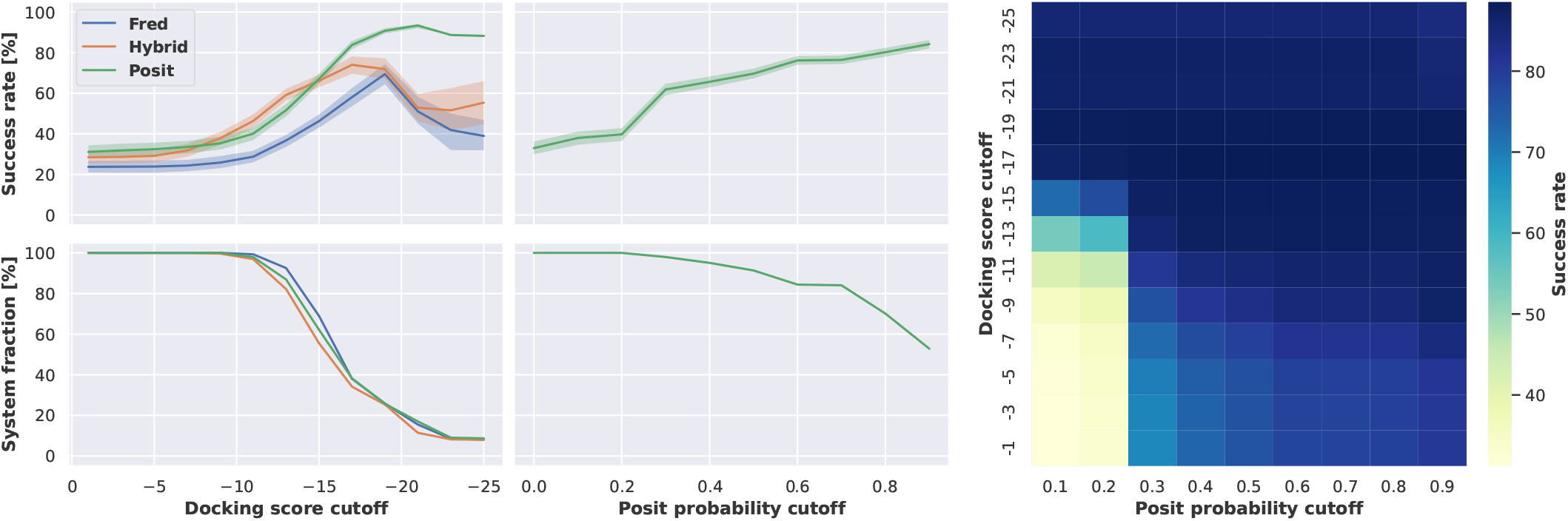
Docking success rates improve when applying more stringent docking score or Posit probability cutoffs. Figure version without normalization of docking score by number of heavy atoms, compare to **Figure 4**. Reported success rates and confidence intervals were estimated using bootstrapping.

**Figure 11.**
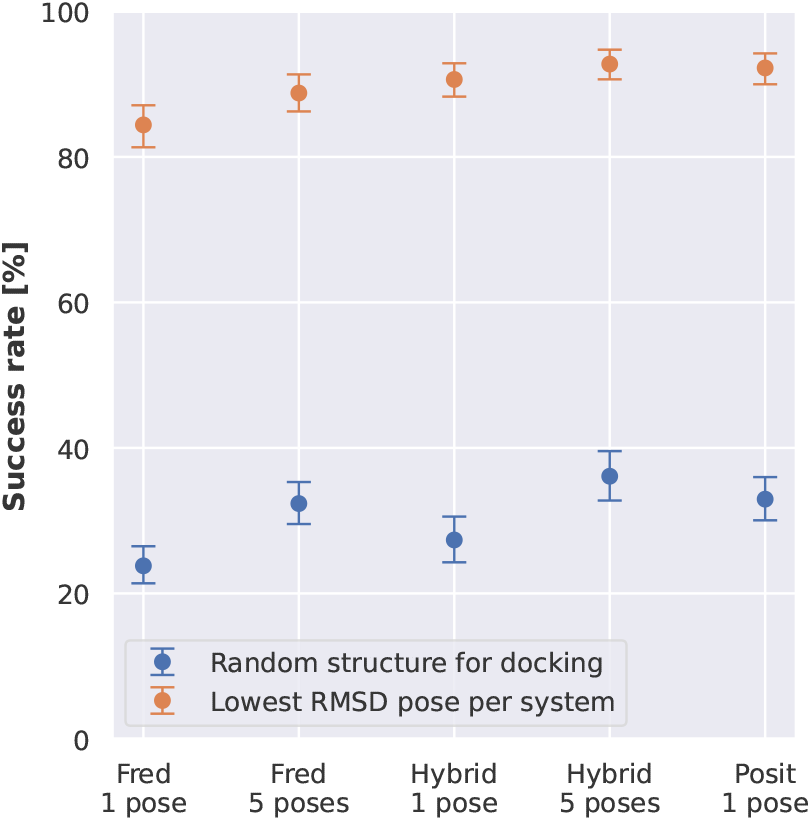
Returning multiple docked poses does significantly improve the success rate of generating a low RMSD docking pose when docking into a single randomly chosen structure. When docking into all available structures returning multiple poses does not significantly improve the best attainable success rate. Posit was not included since the used Openeye Toolkits (2021.1.1) version did not support the generation of multiple poses. Reported success rates and confidence intervals were estimated using bootstrapping.

**Figure 12.**
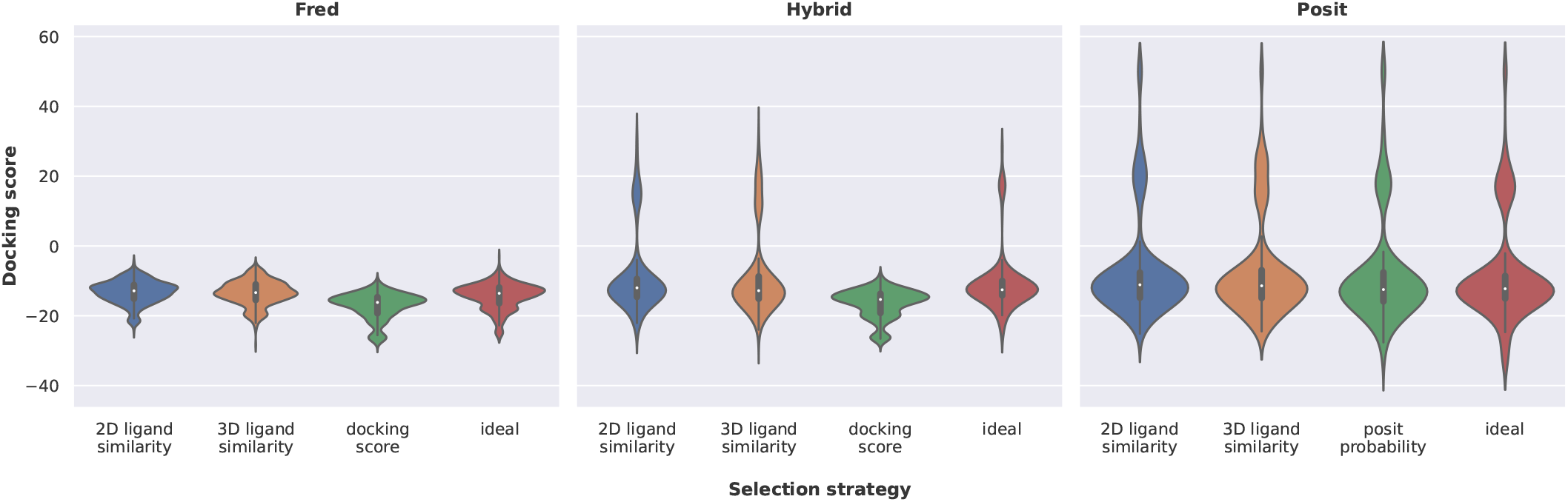
Posit and Hybrid generate more docking poses with high docking score compared to Fred. Poses selected by 2D or 3D ligand similarity to the co-crystallized ligand, by docking score / Posit probability, or by identifying the lowest RMSD pose (ideal scenario) for different docking methods. An upper bound of 50 was introduced for the docking scores to allow proper visualization. Docking with the Fred method and selecting docking poses via the docking scores generates docking score distributions with the lowest docking scores. Docking with Posit generates several high docking score poses that indicate atom clashes which may require additional structural optimization.

**Figure 13.**
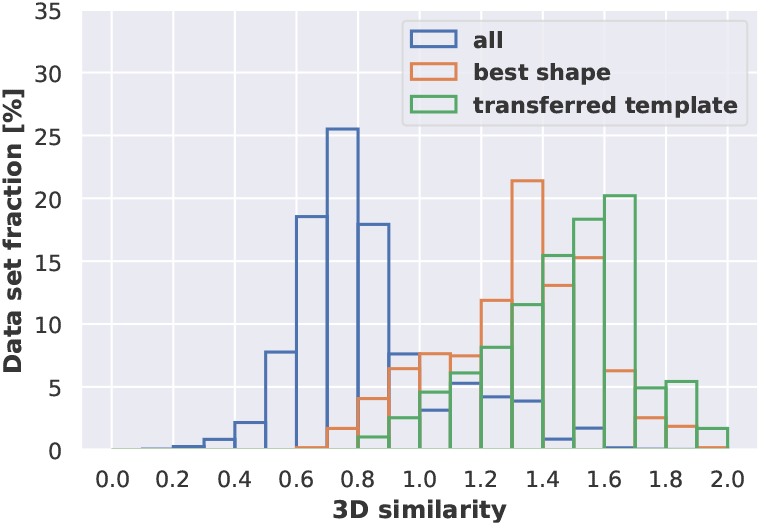
Comparison of Posit ligand template similarity for different docking experiments. When considering all ∼40K docking runs, the average ligand similarity according to shape and electrostatics between co-crystallized ligand and molecule to dock was below 1. Restricting this analysis to the structure with the most similar co-crystallized ligand increased the average 3D similarity, which has a beneficial effect on the docking performance for all tested docking methods (**Figure 5**). This 3D similarity to the co-crystallized ligands can be even more increased when transferring more similar ligands ligands from other kinases. However, this strategy only was found to improve performance when docking into a single randomly selected kinase structure (**Figure 6**).

The human AurA structures included in this benchmark data set were crystallized in the DFG in/αC helix in conformation and the DFG out-like/αC helix in conformation, which represents a relatively small conformational change. The docking success rate did not significantly differ when comparing the results of docking into structures of the same kinase conformation only or in structures of both kinase conformations (**Figure 7** left). Also, the structure with the best Posit probability was in the majority (76.4 %) of docked ligands in the correct kinase conformation (**Figure 7** middle).

Human BRAF was crystallized in the DFG in/αC helix out conformation and the DFG out/αC helix in conformation, which represents a rather large conformational change. Again, the docking success rate did not differ significantly when comparing the results of docking into structures of the respective kinase conformation only or in structures of both kinase conformations (**Figure 7** left). However, compared to the relatively small conformational change in case of human AurA, the Posit method was able to select the correct kinase conformation for almost all ligands docked (96.9 %, (**Figure 7** middle)).

These results show that Posit and potentially other docking tools are more likely to identify the correct kinase conformation in case of larger structural rearrangements. However, due to the flexible nature of kinases, it is possible that structures selected by docking that our experimental approach annotates as putative incorrect kinase conformations (in the case of human AurA) may actually represent a viable bound conformation. Such structure may simply have not been observed crystallographically due to crystallization conditions (**Table 2**) or crystal packing artifacts [38]. In aqueous solution, more than one conformation may be important for ligand binding even if not observed in an X-ray structure [32].

## Conclusions

Employing protein:ligand complex information for predicting bioactivity with ML models holds significant potential to improve performance [7, 10, 11]. Combining such models with highly automated and accurate docking pipelines is both critical for predicting the bioactivity of newly designed compounds and can be useful for enlarging the dataset available for training by providing structures for training data for which only activities are available [39].

In this study, we presented a cross-docking benchmark for highly automated docking pipelines integrated into the [KinoML](https://github.com/openkinome/kinoml) framework. The benchmark data set focuses on protein kinases and was carefully curated with regard to the flexible nature of kinases and comprises of 589 protein kinases co-crystallized with 423 different ATP-competitive ligands.

We found that the investigated docking methods Fred, Hybrid, and Posit from the OpenEye Toolkits [34] were able to recover the majority of ligand poses observed in the experimentally resolved structures. However, the docking performance was dependent on the docking method and the selected kinase structure for docking. Methods biased by the co-crystallized ligand (Hybrid and Posit) were more successful than the standard docking method Fred. Docking into kinase structure with more similar co-crystallized ligand was found to positively impact docking performance, which is in line with previous cross-docking studies [19, 20, 21]. Also, better docking scores and Posit probabilities indicated lower RMSD docking poses. However, such approach requires docking into multiple structures, whereas the ligand similarity to the co-crystallized ligand can be calculated before performing the more computational expensive docking calculation. In addition, docking into multiple structure strongly increased the chance to generate a low RMSD docking pose for all investigated docking methods, e.g., from 33.1 % when docking into one random struc-ture with Posit to 83.2 % when docking into 20 random structures (**Figure 2** right)). Interestingly, the docking performance of the most successful docking method Posit was not affected by docking into different kinase conformations. And the Posit probability was able to predict the correct kinase conformation in the majority of tested kinase systems.

This benchmark was not only performed to test the designed docking pipelines but also to learn for future structure-informed machine learning experiments for bioactivity prediction. Based on our findings, the Posit docking method was the most successful to generate docking poses with the lowest RMSD when docking into the kinase structure with the most similar co-crystallized ligand. If the downstream ML model is able to handle multiple input docking poses for a single kinase:inhibitor pair, the model could learn the relevance for each input docking pose based on ligand similarity and docking score. Also, the ML model is more likely to see a low RMSD docking pose when providing poses from docking into multiple structures rather than multiple poses from docking into the same structure.

Finally, it would be interesting to see the performance of the newly developed ML-based docking approaches in concert with bioactivity predictions for this benchmark data set [12, 13, 14], but this experiment was not included in this study.

## Data availability

The kinase-docking-benchmark repository is made publicly available and stores all relevant Python scripts and Jupyter notebooks to reproduce the presented docking benchmark results and analysis. The associated docking pipelines are available in the [KinoML](https://github.com/openkinome/kinoml) framework. The generated docking poses are stored in an [OSF project](https://osf.io/f6tgm/).

## Acknowledgments

The authors thank OpenEye Scientific Software for providing a free academic license to the OpenEye Toolkits to the Chodera lab.

## Detailed Methods

### Benchmark data set

The OpenCADD-KLIFS module [26, 27] was used to query for available kinase structures in the PDB [25] in January 2022 (**Figure 3**). The retrieved 12, 572 KLIFS structures were filtered for the presence of a single ligand in the ATP binding site, since the docking methods Hybrid and Posit require a co-crystallized ligand for pocket definition and the presence of multiple ligands in the same binding pocket likely hinders correct sampling of the docking programs. For each PDB entry, only the highest quality chain was retained according to the structure quality reported by KLIFS leading to 5, 017 remaining entries. Next, KLIFS entries were excluded that contain mutations in the 85 KLIFS residues defining the binding site as well as KLIFS entries with an incompletely resolved ATP binding site. This step should assure that modeling of missing or mutated residues does not bias the docking benchmark and left 1, 914 KLIFS entries for further processing. The remaining entries were filtered for a reasonable molecular weight (150 1, 000 Da) of the co-crystallized ligand, and for SMILES representation stored in the PDB interpretable by [RDKit](http://www.rdkit.org/) and OpenEye Toolkits [34] that are later used for docking and analysis. For example PDB entry 5C01 does not have a valid SMILES representation stored in the PDB. This step removed 27 KLIFS entries. Next, structures were removed with multiple instances of the same ligand bound to the same protein chain removing another 27 KLIFS entries. The remaining 1, 888 KLIFS entries were grouped to select those kinases that have at least 10 different KLIFS entries available. Finally, the most populated kinases were picked for each kinase group (TK, TKL, STE, …) and conformation (DFG/αC helix in/in, in/out, outlike/in and out/in) to assure covering different kinase groups and conformations leaving the final benchmark data set comprising 589 KLIFS entries co-crystallized with 423 different ligands (**Table 1**).

### Molecular docking

Protein structure preparation was performed using OEChem and Spruce functionality from the OpenEye Toolkits 2021.1.1 [34] implemented into the [KinoML framework](https://github.com/openkinome/kinoml/commit/e2bead16cb8e03c89ec7af0f5e0167b81adbe03e). Missing side chains were built using the Spruce toolkit, and all protein chain termini capped with NME and ACE residues. The resulting protein:ligand complex was protonated at pH 7.4 using the OEChem toolkit.

Molecular docking was performed using OEDocking, Omega and Quacpac functionality from the OpenEye Toolkits 2021.1.1 [34] implemented into the [KinoML framework](https://github.com/openkinome/kinoml/commit/e2bead16cb8e03c89ec7af0f5e0167b81adbe03e).

For each molecule to dock reasonable tautomers at pH 7.4 were determined using Quacpac. Undefined stereo centers were enumerated with Omega. Finally, up to 800 conformations were generated for each enantiomer and tautomer using Omega with sampling in Pose mode.

For the Fred and Hybrid docking algorithms, docking was performed with high search resolution. The 5 best docking poses according to the Chemgauss4 docking score were returned and written to disk. Posit docking calculations were performed with the IgnoreNitrogenStereo option set to True, which is the recommended setting according to the [documentation](https://docs.eyesopen.com/toolkits/python/dockingtk/poseprediction.html?highlight=setignorenitrogenstereo). The pose relaxation was turned off and the posed molecule subsequently scored with the Chemgauss4 docking score.

### Docking pose analysis

For calculation of the heavy atom RMSD of a docking pose to its reference X-ray structure, a structural superposition was performed to align kinase binding site residues, as described below. This step involved Spruce functionality from the OpenEye Toolkits 2021.1.1 [34] implemented into the [KinoML framework](https://github.com/openkinome/kinoml/commit/e2bead16cb8e03c89ec7af0f5e0167b81adbe03e). The superposition was thereby restricted to the heavy atoms of the 85 KLIFS residues retrieved via the OpenCADD-KLIFS module [26, 27]. Subsequently, the heavy atom RMSD of co-crystallized ligand and docking pose was calculated using SpyRMSD [40].

The 2D similarity between the small molecule to be docked and the co-crystallized ligand was calculated with Morgan fingerprints with features, radius of 2 and 2, 048 bits implemented in the [RDKit](http://www.rdkit.org/) library. The 3D similarity in form of shape and electrostatics was determined for enumerated conformations of the small molecule to dock to the 3D structure of the co-crystallized ligand. The overlay was performed using Shape, Omega, and Quacpac functionality from the OpenEye Toolkits 2021.1.1 [34] implemented into the [KinoML](https://github.com/openkinome/kinoml/commit/e2bead16cb8e03c89ec7af0f5e0167b81adbe03e) framework. For each molecule reasonable tautomers at pH 7.4 were determined using Quacpac. Undefinded stereo centers were enumerated with Omega. Finally, up to 200 conformations were generated for each enantiomer and tautomer using Omega with sampling in Classic mode. These conformations were overlayed to the 3D structure of the co-crystallized ligand and the 3D similarity determined using the TanimotoCombo score considering shape and electrostatics.

### Statistical measures

Per experiment, 40*K* docking runs were performed in this cross-docking study docking each ligand into each structure of the same kinase and conformation. To estimate the impact of, e.g., ligand similarity or docking score on docking performance the success rates including 95 % confidence intervals were calculated using 1, 000 bootstrap samples.

For example, the reported success rate of 33.0 % for Posit when docking a ligand into a randomly selected structure of the same kinase and conformation (**Figure 2**) was generated by (i) randomly picking a single docking pose for each ligand, (ii) calculating the mean success rate over all structures, (iii) performing step i and ii 1000 times and (iv) calculating the mean success rate over all bootstrap samples as well as the 95 % confidence interval.

The detailed procedure for each analysis can be found in the jupyter notebooks of the accompanying kinase-docking-benchmark repository.

## Appendix

